# Concurrent tACS-fMRI reveals causal influence of power synchronized neural activity on resting state fMRI connectivity

**DOI:** 10.1101/122820

**Authors:** Marc Bächinger, Valerio Zerbi, Marius Moisa, Rafael Polania, Quanying Liu, Dante Mantini, Christian Ruff, Nicole Wenderoth

**Affiliations:** Neural Control of Movement Lab, Dept. of Health Sciences and Technology, ETH Zürich, 8057 Zürich, Switzerland; Laboratory for Social and Neural Systems Research (SNS), Dept. of Economics, University of Zürich, 8006 Zürich, Switzerland; Movement Control and Neuroplasticity Research Group, Department of Kinesiology, KU Leuven, Belgium; Institute for Biomedical Engineering, University and ETH of Zurich, 8052 Zürich Switzerland; Neuroscience Centre Zurich (ZNZ), Zurich, Switzerland

## Abstract

Resting state fMRI (rs-fMRI) is commonly used to study the brain’s intrinsic neural coupling, which reveals specific spatiotemporal patterns in the form of resting state networks (RSN). It has been hypothesized that slow rs-fMRI oscillations (<0.1 Hz) are driven by underlying electrophysiological rhythms that typically occur at much faster timescales (>5 Hz); however, causal evidence for this relationship is currently lacking. Here we measured rs-fMRI in humans while applying transcranial alternating current stimulation (tACS) to entrain brain rhythms in left and right sensorimotor cortices.

The two driving tACS signals were tailored to the individual’s alpha rhythm (8-12 Hz) and fluctuated in amplitude according to a 1 Hz power envelope. We entrained the left versus right hemisphere in accordance to two different coupling modes where either alpha oscillations were synchronized between hemispheres (phase-synchronized tACS) or the slower oscillating power envelopes (power-synchronized tACS).

Power-synchronized tACS significantly increased rs-fMRI connectivity within the stimulated RSN compared to phase-synchronized or no tACS. This effect outlasted the stimulation period and tended to be more effective in individuals who exhibited a naturally weak interhemispheric coupling. Using this novel approach, our data provide causal evidence that synchronized power fluctuations contribute to the formation of fMRI-based RSNs. Moreover, our findings demonstrate that the brain’s intrinsic coupling at rest can be selectively modulated by choosing appropriate tACS signals, which could lead to new interventions for patients with altered rs-fMRI connectivity.

**Significance Statement:** Resting state fMRI has become an important tool to estimate brain connectivity. However, relatively little is known about how slow hemodynamic oscillations measured with fMRI relate to electrophysiological processes.

It was suggested that slowly fluctuating power envelopes of electrophysiological signals synchronize across brain areas and that the topography of this activity is spatially correlated to resting state networks derived from rs-fMRI. Here we take a novel approach to address this problem and establish a causal link between the power fluctuations of electrophysiological signals and rs-fMRI via a new neuromodulation paradigm, which exploits these power-synchronization mechanisms.

These novel mechanistic insights bridge different scientific domains and are of broad interest to researchers in the fields of Medical Imaging, Neuroscience, Physiology and Psychology.

## Introduction

Resting state functional magnetic resonance imaging (rs-fMRI) is a widely-used tool for investigating large-scale functional connectivity within the human brain (Biswal et al., 1995; Fox and Raichle, 2007). Rs-fMRI measures spontaneous slow fluctuations (i.e. <0.1 Hz) of the blood-oxygen level dependent (BOLD) signal at rest. These fluctuations form spatial patterns of correlated activity that can be mapped onto resting state networks (RSN); these exhibit a unique topography that often resembles networks activated by specific tasks (i.e. sensorimotor network, visual network, etc.) (Damoiseaux et al., 2006). Alterations in resting-state connectivity have been associated with several neuropathologies (Greicius, 2008; Bullmore and Sporns, 2009; Fornito and Bullmore, 2010; Alaerts et al., 2014; 2016) opening new opportunities for identifying disease-specific biomarkers and potential therapy targets at the brain circuit level (Alaerts et al., 2014; Woolley et al., 2015). However, which exact electrophysiological mechanisms cause RSNs to emerge is still debated making it difficult to design new interventions that target activity at the cell population level with the aim of normalizing large-scale connectivity within specific circuits. Previous studies have linked rs-fMRI connectivity to ultra-slow (<0.5 Hz) fluctuations of electrophysiological signals (Pan et al., 2013), or to positively correlated power envelopes (also called band-limited power BLP signals) of faster frequency bands including the delta (<4Hz) (Lu et al., 2007), alpha (8-12 Hz)/beta (12-14 Hz)(Mantini et al., 2007; Brookes et al., 2011; Wang et al., 2012) and gamma bands (He et al., 2008; Nir et al., 2008; Schoelvinck et al., 2010). This converging evidence indicates that slowly fluctuating components of the electrophysiological signal (either the power envelope or the signal itself) synchronize across different areas of the brain, and the topography of this synchronization pattern has been linked to heightened BOLD connectivity within corresponding RSNs.

However, these previous studies revealed mainly correlative evidence arguing that the topography or strength of connectivity is similar when RSNs determined from synchronized activity of the BOLD signal were compared to RSNs determined from electrophysiological readouts (LFP or EEG) (Lu et al., 2007; He et al., 2008; Nir et al., 2008; Brookes et al., 2011; Wang et al., 2012; Hipp and Siegel, 2015). Here we used a novel interventional approach to experimentally probe the hypothesis that synchronizing power envelopes of the prominent alpha rhythm across anatomically connected brain areas increases rs-fMRI connectivity within the targeted network. The basic idea is to modulate electrophysiological activity with non-invasive electrical stimulation applying appropriately designed, EEG-based driving signals while measuring the induced changes in rs-fMRI connectivity. This approach is not only interesting as a research tool but also as a potential intervention for modulating large-scale connectivity at rest within clinically relevant circuits.

## Materials & Methods

We applied transcranial alternating current stimulation (tACS) to entrain endogenous neuronal oscillations in a frequency-dependent way (Zaehle et al., 2010; Polanía et al., 2012; Herrmann et al., 2013; Helfrich et al., 2014; Cecere et al., 2015) without causing significant interference with MRI measurements (Antal et al., 2014). Using a three-electrode setup to drive neural activity within left and right sensorimotor cortices (SM1), functional coupling between hemispheres can be either strengthened or weakened by administering two iso-frequent stimulation signals which are either in-phase or anti-phase, respectively (Polanía et al., 2012; 2015). The waveforms were tailored to the subject’s individual alpha frequency band (IAF; corresponding roughly to 8-12 Hz oscillation), which was chosen as a “carrier frequency” because it is a strong endogenous rhythm present at rest. Moreover, in motor cortex the power envelope of the alpha band has been linked to interhemispheric connectivity measured by EEG/MEG (Siems et al., 2016), rs-fMRI connectivity (Mantini et al., 2007) and task-based fMRI activity (Ritter et al., 2009). Importantly, we modulated the amplitude of the tACS signals according to a 1 Hz envelope, thereby mimicking a cross-frequency coupling phenomenon (i.e., a phase-amplitude coupling mechanism between the alpha and delta rhythm) that emerges in humans at rest (Siems et al., 2016). We then aimed to disambiguate whether interhemispheric rs-fMRI connectivity is mainly driven by the interhemispheric synchronization of 1 Hz amplitude envelopes, or by the synchronization of left and right IAF. We therefore stimulated SM1 of each hemisphere with the same amplitude-modulated IAF driving signal, but with different phase relationships between hemispheres. In the *power-synchronized tACS* condition, the 1Hz amplitude envelopes were in-phase but the IAF signals were anti-phase; by contrast, in the *phase-synchronized tACS* condition, the IAF signals were in-phase but the 1 Hz amplitude envelopes were anti-phase. Importantly, the power-synchronized tACS condition approximates the interhemispheric coupling as measured beforehand with EEG at rest, whereas the phase-synchronized tACS inverts this interhemispheric phase–relationship and acts as negative control. We hypothesized that the power-synchronized stimulation regime would increase rs-fMRI connectivity between hemispheres.

### Experimental Design

The experiment consisted of two sessions, which were on average 1 week apart. During the first session 5 minutes of resting-state EEG (eyes-open) was measured and the participants’ EEG frequency spectrum was pre-screened. Only subjects with a clear alpha-peak participated in the tACS/fMRI experiment. Based on the prescreening, we tailored the tACS stimulation signals to the individual alpha-frequency (IAF). In the second session, a combined tACS/resting-state fMRI (eyes-open) experiment was conducted to test the effects of two different stimulation paradigms on rs-fMRI connectivity between the sensorimotor cortices of the two hemispheres.

### Subjects

35 subjects were recruited for the EEG session. All subjects provided informed consent as approved by the Research Ethics Committee of the Canton of Zurich. Subjects were pre-screened to have a clear alpha-peak over sensorimotor areas (at electrode C3 or C4; see EEG analysis for definition of alpha-peak). 22 subjects showed an alpha peak over both hemispheres. Their EEG was analyzed in more detail and they participated in the subsequent tACS/fMRI experiment. From the 22 subjects who were scanned, two were excluded from analysis; one because of too much movement and one fell asleep during the experiment. 20 subjects were analyzed and all reported results (EEG and fMRI) are from those subjects unless stated otherwise (n = 20, age = 24.8 ± 4.1 years, mean ± s.d., 10 females, 19 righthanded by self-indication).

### EEG acquisition

Electronecephalography (EEG) was measured using a 128-channel HydroCel Geodesic Sensor Net (GSN) using Ag/AgCl electrodes provided by Electrical Geodesics (EGI, Eugene, Oregon, USA). This system uses the vertex (Cz) electrode as physical reference. EEG recordings, electrooculograms for horizontal and vertical eye movements respectively, and an electromyogram for the muscular noise associated with swallowing were recorded in parallel with a sampling frequency of 1000 Hz.

During EEG acquisition subjects sat in a dark room and fixated on a cross presented on a computer screen in front of them for five minutes. The unfiltered data were saved for offline analyses.

### EEG preprocessing

EEG signals were bandpass filtered off-line (2-40 Hz) and further processed using independent component analysis (ICA) for the removal of ocular and muscular artifacts with eeglab (Delorme and Makeig, 2004). After ICA decomposition the artifact ICs were automatically detected by correlating their power time-courses with the power time courses of the electric reference signals: the horizontal electrooculogram (hEOG), the vertical electrooculogram (vEOG) and electromyogram (EMG) at the base of the neck. The data was then downsampled to 200Hz and re-referenced to the common average (Liu et al., 2015) to remove the bias towards the physical reference site (Luck, 2014).

### EEG source localisation

After the fMRI experiment (see below) source localization of the EEG data was performed to control that the results obtained in sensor space indeed reflected activity in sensorimotor areas. A forward head model was built with the finite element method (FEM), using a 12-tissue head template and the standard electrode positions for a 128-channel EGI cap. The head template was obtained from the IT IS foundation of ETH Zurich (Iacono et al., 2015) and included 12-tissue classes (skin, eyes, muscle, fat, spongy bone, compact bone, cortical gray matter, cerebellar gray matter, cortical white matter, cerebellar white matter, cerebrospinal fluid and brain stem). Specific conductivity values were associated with each tissue class (i.e. skin 04348 S/m, compact bone 0.0063 S/m, spongy bone 0.0400 S/m, CSF 1.5385 S/m, cortical gray matter 0.3333 S/m, cerebellar gray matter 0.2564 S/m, cortical white matter 0.1429 S/m cerebellar white matter 0.1099 S/m, brainstem 0.1538 S/m, eyes 0.5000 S/m, muscle 0.1000 S/m, fat 0.0400 S/m (Haueisen et al., 1997). The dipoles corresponding to brain sources were placed on a regular 6-mm grid spanning cortical and cerebellar gray matter. After the head model template was established, the brain activity in each dipole source was estimated by the exact low-resolution brain electromagnetic tomography (eLORETA (Pascual-Marqui et al., 2011)) for each subject.

### EEG analysis

EEG analysis was performed using the fieldtrip toolbox (Oostenveld et al., 2011) and custom Matlab scripts. Group analyses in sensor space were verified by parallel analyses in source space while individual-difference analyses were restricted to sensor space to minimize confounds caused by imperfect source localization. For each subject the individual alpha-peak and the correlational properties of the alpha rhythm during rest was determined for electrodes C3 and C4 (sensor space), which corresponds to the left and right primary sensorimotor cortices, respectively.

For determining the individual alpha peak we first calculated the frequency spectrum for each electrode using the absolute of the Fourier transform. The individual alpha peak was determined as the local maximum between 7 and 14 Hz of the resulting spectrum. Only subjects with a clearly identifiable alpha peak in sensor space when visually inspected were included in the fMRI experiment. The individual alpha frequency band (IAF) for each subject was then defined as individual alpha-peak +/− 2 Hz (Zaehle et al., 2010). We then bandpass filtered the signal within the IAF of each subject (IAF signal). We also calculated the amplitude envelopes of the IAF signals using the absolute of the Hilbert transform. Next, the natural phase relationship between the two IAF signals and the two amplitude envelopes of C3 and C4 was determined via the Pearson’s correlation coefficient r and via the instantaneous phase relationship. The latter was calculated by epoching the resting-state EEG data into segments of 5 seconds. For each segment we calculated the frequency spectrum, the instantaneous phase of the IAF signal and the instantaneous phase of the amplitude envelope. We averaged the frequency spectrum across all 5 second segments and determined mean IAF peaks and 95% confidence intervals. The plots were visually inspected for each individual to confirm that the IAF exhibited a clear peak in the 5 sec intervals to ensure that the relative phase would represent meaningful values (Figure 1A). Instantaneous phase difference of the IAF signal and the amplitude envelope between C3 and C4 was calculated for each subject and the main directionality of the phase difference across subjects was statistically assessed using V tests for circular uniformity. In short V tests assess whether a vector of angles has a known main direction(Berens, 2009). We tested 180° for the IAF signal and 0° for the amplitude envelopes. The same phase analysis was then performed in source space, i.e. for the first principal component of the signals extracted from the center of gravity (cog) of the left and right part of the sensorimotor network as identified during the fMRI analysis (cog left hemisphere: MNI −15 −29 63, cog right hemisphere: MNI 13 −28 63) This was done to confirm that the signals measured in C3/C4 originated from the sensorimotor network.

**Figure 1.**
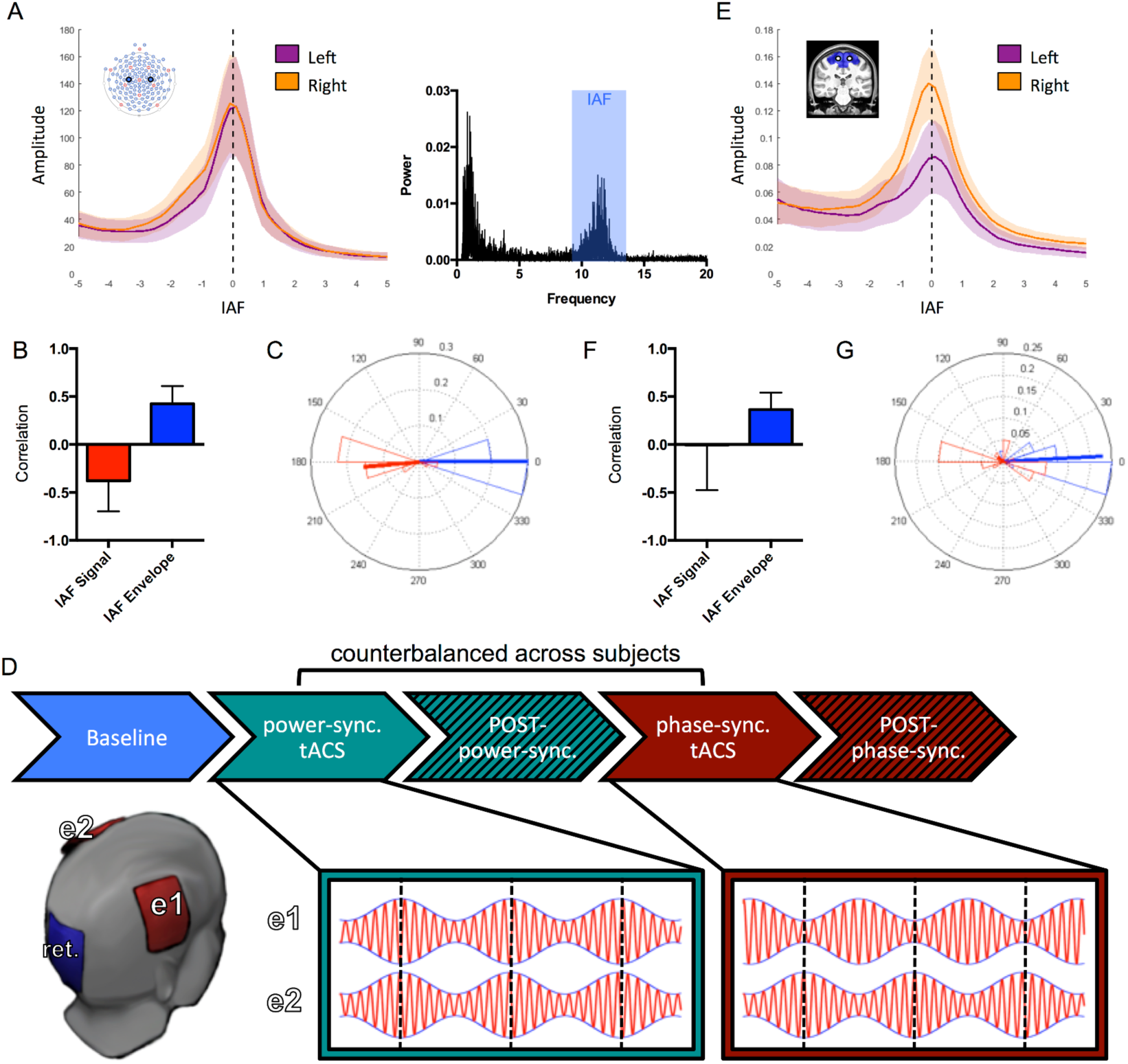
EEG-based stimulation signals during fMRI. (A) For each subject the individual alpha peak and individual alpha frequency band (IAF = alpha peak +/− 2Hz) was determined from electrodes C3/C4 during rest (eyes-open). The right panel shows an exemplary frequency spectrum. The left panel shows the mean and 95% confidence interval (shaded area) of the EEG frequency spectrum normalized to the IAF +/−5 Hz of the left and the right hemisphere, indicating a clear peak. (B) Correlational analysis showing an anti-correlated relationship between electrodes C3/C4 for the signal within IAF (red) and a positive correlation for the envelope modulation of IAF (blue). (C) Histogram of relative phase difference between C3 and C4 of IAF signal (red) and envelope (blue) across subjects. Arrows representing average phase difference across subjects (D) Based on the subject’s IAF, two kinds of stimulation signals were applied via a 3-electrode setup during resting-state fMRI. Power-synchronized tACS mimics the in-phase relationship of the IAF power envelopes between C3/C4 (while the IAF signal was anti-correlated; blue-green) and phase-synchronized tACS (control) with the opposite correlational properties (antiphase power-fluctuations, synchronized IAF signal). 35 minutes of resting state fMRI were recorded, split into 5 runs of 7 minutes from the same 20 subjects as in the EEG experiment (see methods). During runs 2 and 4, power-synchronized and phase-synchronized tACS was applied via 2 active electrodes placed on the left and right motor cortex (red), run 1 served as baseline for further analysis and runs 3 and 5 measured potential after-effects. (E) Post-fMRI analysis of the EEG data confirms that the IAF is clearly detectable in source space, i.e. in left and right sensorimotor cortex. (F,E) Correlational and instantaneous phase properties in source space. Note that the IAF envelope exhibits robust synchronization (i.e. 0 deg relative phase) between hemispheres while the IAF signals exhibt large inter-subject variability.

### tACS 3-electrode setup

We used two modified MR compatible DC stimulator plus devices (Neuroconn GmbH, Illmenau, Germany) that were connected to two active electrodes and one common return electrode (electrode-size 5×7 cm). The active electrodes were placed over the left and right motor cortex as determined by transcranial magnetic stimulation (hotspot of the first dorsal interosseus muscle) and the return electrode was placed approximately 2-3 cm above the inion to minimize the perception of phosphenes while preventing that subjects would not lie on the return electrode while scanning (which might have caused discomfort).

The stimulation was controlled via the REMOTE connectors of the stimulators which allows externally generated voltage signals to be translated into current/stimulation signals. The actual stimulation signals were produced with a custom-made Matlab script and sent to the stimulators via a National Instruments Card (NI-USB 6343). This allowed precise control of the output signals (especially phase stability between the two signals).

The stimulation signals were individualized to the subject’s IAF. If the individual peak frequency differed between left and right hemispheres then the average of the two was used. Importantly since the precise frequency of the individual alpha-peak might change slightly from the EEG session to the fMRI session (cmp. (Vossen et al., 2014)), we used stimulation signals that covered the whole IAF band (alpha peak +/−2 Hz). The signals were based on the subject’s IAF that was modulated with a 1 Hz envelope (Figure 1D). The 1 Hz envelope was chosen based on pilot data where the group average frequency spectrum had a broad peak from 0.5 to 2 Hz with a maximum at 1 Hz. We used this value for designing the tACS waveforms used in the main experiment. We repeated this analysis with the full sample, which revealed a group maximum at 1.2 Hz. Note that all these analyses were performed after applying a bandpass filter to the EEG signal with cutoffs at the IAF +/− 2Hz. The maximum current intensity (peak-to-peak amplitude) over the active electrode was1.5 mA and 0.75 mA during the peak and trough of the envelope, respectively.

Two different phase relationships between the left and right hemisphere were tested for the experiment. During power-synchronized tACS the two stimulation signals were in anti-phase (180° degrees phase-shifted) and the envelopes were in-phase (0° phase shifted). During phase-synchronized tACS the stimulation signals were inphase (0° phase shifted) and the envelopes were anti-phase (180° phase-shifted). Note that power-synchronized tACS represents a synchronization of the power envelopes similar to that observed for the EEG signals measured over C3 and C4, whereas phase-synchronized tACS inverts this phase relationship (i.e. no synchronization of the power envelope), and is used as negative control condition.

### MRI acquisition

Resting-state imaging was performed at the Laboratory for Social and Neural Systems research (SNS-Lab) of the University of Zurich, on a Philips Achieva 3T whole-body scanner equipped with an eight-channel MR head coil. Before applying tACS during fMRI we did basic safety and quality tests for the three-electrode setup. We tested for dynamic tACS artifacts and for heating under the tACS electrodes, following the protocol described previously in (Moisa et al., 2016). In short, the conducted analyses did not reveal dynamic artifacts due to our tACS stimulation or an increase in temperature due to up to 30 min of stimulation. Moreover, data-driven artifact removal performed with FSL-FIX (see rs-fMRI analysis) did not reveal any components with a topography that would suggest tACS-related artifacts (for example we did not see components under/close to the electrodes).

Initially, T1-weighted 3D turbo field echo B0 scans were acquired for correction of possible static distortion produced by the presence of the active electrode (voxel size = 3×3×3 mm^3^, 0.5 mm gap, matrix size = 80 × 80, TR/TE1/TE2 = 418/4.3/7.4 ms, flip angle = 44, no parallel imaging, 37 slices). High-resolution T1-weighted 3D turbo field echo structural scans were acquired and used for image registration and normalization (181 sagittal slices, matrix size = 256 × 256, voxel size = 1 mm^3^, TR/TE/TI = 8.3/2.26/181 ms). Thereafter, five resting-state runs of 7 minutes were collected for each subject (2-3 minutes break between each block). Each resting-state run contains 200 volumes (voxel size = 3×3×3 mm^3^, 0.5 mm gap, matrix size = 80 × 80, TR/TE = 2100/30 ms, flip angle = 79, parallel imaging factor =1.5, 35 slices acquired in ascending order for full coverage of the brain).

The first run served as a baseline measurement. During the second and fourth runs tACS was applied (the order of tACS conditions was counterbalanced between subjects). Runs 3 and 5 captured potential after-effects (Figure 1). This allowed us to compare the effect of each stimulation protocol on rs-fMRI connectivity in relation to baseline, but also to directly contrast the effects of power-synchronized vs. phase-synchronized tACS stimulation.

### rs-fMRI analysis

FMRI analysis was performed similar to the protocol previously used by Stagg et al. (2014). Brain functional networks (i.e. independent components) were identified using the Multivariate Exploratory Linear Optimized Decomposition into Independent Components (MELODIC; version 3.10) module in FSL (FMRIB’s Software Library, www.fmrib.ox.ac.uk/fsl). Standard pre-processing consisted of motion correction, brain extraction, spatial smoothing using a Gaussian kernel of FWHM 8.0 mm and high-pass temporal filtering (100 s / 0.01 Hz). Additionally, artifact components from non-neural sources were removed with FSLs FMRIB’s ICA-based Xnoiseifier (FSL-FIX (Griffanti et al., 2014; Salimi-Khorshidi et al., 2014)). In short FSL-FIX automatically classifies each component into either signal or noise, and regresses the noise components from the original data using the standard classifier at threshold 20. In comparison with a manual classification done by two experts on a subset of the subjects (n=5), FSL-FIX yielded a very high sensitivity (97.6%) in detecting true RSNs.

The artifact-cleaned functional data were then aligned to structural images and normalized into MNI space using linear and non-linear transformations (ANTs, advanced normalization tools, http://stnava.github.io/ANTs).

Normalized functional data for each subject were temporally concatenated across subjects to create a single 4D dataset and group ICA was performed to identify resting state networks (RSNs). Between-subject analysis was performed using a dual regression approach implemented in FSL (Beckmann et al., 2009). In short this approach consists of two stages: (1) A spatial regression of the data is calculated to identify the timecourse of a RSN. (2) A temporal regression with those timecourses is determined to get the subject specific map of the RSN. This is done for each RSN.

The resulting subject-specific component map was then masked by the 75^th^ percentile group mean RSN map. The mean value of the parameter estimates within this region was extracted for each subject. The average of this parameter estimate can be seen as a measure of the average strength of functional connectivity within each RSN. This analysis was performed for each RSN separately.

The average scores of each subject were then submitted to a linear mixed effects model (LMEM) with the fixed factor stimulation type (5-levels, repeated measure) and the random factor subject. Corresponding contrasts were used for post-hoc pairwise comparisons (LSD) for which effect sizes were calculated using Cohen’s d as an intuitive estimate. This was done for each RSN of interest separately.

Finally, a qualitative voxelwise analysis was performed for each RSN using non-parametric paired t-tests (threshold free cluster enhancement; tfce corrected) as implemented by randomise in FSL (Winkler et al., 2014).

### Modelling of the electric field induced by tACS

We compared the electric field induced by phase-synchronized versus power-synchronized tACS using SimNIBS 2.0 (Thielscher et al., 2015) to model the effect. A realistic finite element head model was used for the simulations. We simulated the same electrode placement as during the experiment (i.e. above the left and right hand knob region in M1 and the return electrode 2 cm above the inion). The stimulation signals were downsampled to 80 Hz and a whole brain image of the electric field distribution was calculated for each time point. For visualisation purposes we extracted the norm of the electric field during peaks and troughs of the envelopes of power-synchronized and phase-synchronized stimulation. Furthermore, we investigated how the field fluctuated within our target area, i.e. the sensorimotor network by reconstructing the signal within a 4mm circular seed region at the peak of the electric field during power-synchronized stimulation (MNI: −18, −40, 66). This was done by taking the first principal component of the electric field and upsampling the signal to the original sampling rate of 1280 Hz for visualization purposes (Figure 2).

**Figure 2.**
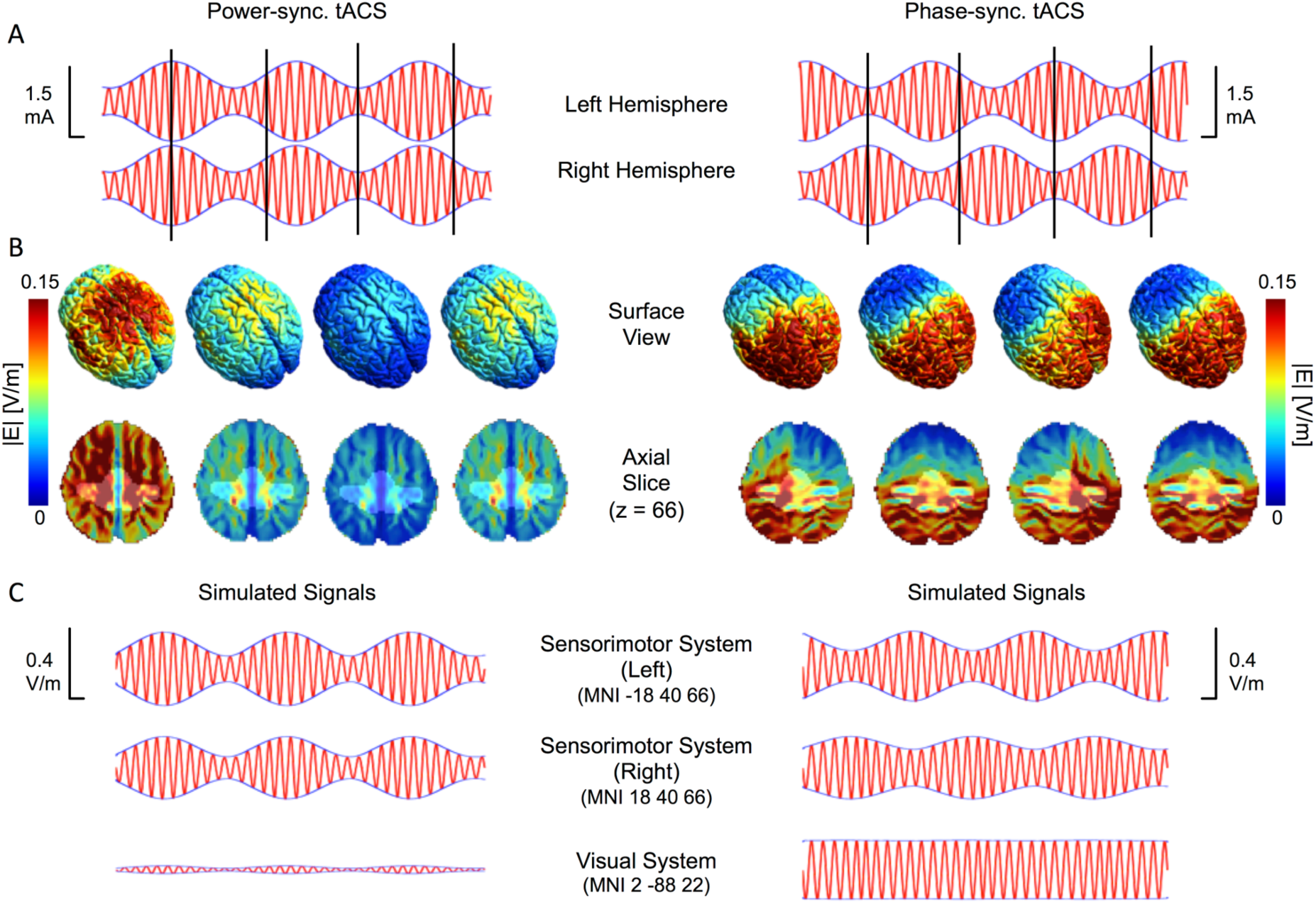
Simulations of the electric field during power-synchronized (left side) and phase-synchronized tACS (right side). (A) The upper part shows the externally applied signals over left and right sensorimotor cortex. (B) The electric field distribution estimated for four different timepoints of the envelope amplitude of the right hemisphere (indicted by black vertical lines) is shown on a surface view and an axial slice through the sensorimotor system (z=66). The highlighted aspect shows the sensorimotor RSN. (C) Simulated signals showing effective modulation within left and right sensorimotor cortex extracted from MNI coordinates +/− 18, −40, 66 (i.e. the maximum of the simulated electric field during power-synchronized stimulation) and the visual cortex (MNI 2, −88, 22; i.e. approximately under the return electrode). Note that during power-synchronized tACS (B,C left) both hemispheres oscillate synchronously between strong and weak stimulation. By contrast, during phase-synchronized tACS (B,C right) one hemisphere is always more strongly activated than the other. Also the maximum modulation depth of the envelope reached similar levels in the sensorimotor cortex for both stimulation conditions (0.25 V/m for power-synchronized tACS and 0.21 V/m for phase-synchronized tACS)

## Results

### Designing EEG-based tACS signals

We recorded and analyzed resting-state EEG in 20 subjects and identified their individual alpha frequency band (IAF, 2 Hz below and above the individual alpha peak, see methods section for inclusion criteria) over left and right SM1 (sensor space, electrodes C3 & C4, respectively; Figure 1A) (Zaehle et al., 2010).

The IAF signals (peak alpha frequency was 10.6 ± 1.2 Hz with an average amplitude of 0.22 ± 0.11 uV over the left hemisphere and 10.5 ± 1.1 Hz with an average amplitude of 0.21 ± 0.13 uV over right hemisphere; all values mean ± s.d) measured in C3/C4 were on average negatively correlated (r_IAF_ = −0.38 ± 0.32) and exhibited an instantaneous relative phase Φ_IAF_ = −174.8° ± 45.2° indicating that the IAF signals of the left and right hemisphere oscillated in anti-phase (Figure 1C red symbols). Next we characterized the IAF envelope (within a 0.1 - 2 Hz band) which was positively correlated between hemispheres (r_Env_ = 0.42 ± 0.19), and tended to oscillate inphase (Φ_Env_ = 0.02° ± 0.21°, Figure 1B,C blue symbols). Using the sensorimotor network from the fMRI analysis (see below) we checked whether the correlational and instantaneous phase properties were still present in source space (Figure 1E,F,G). The peak alpha frequency was comparable to the values obtained in sensor space (left hemisphere: 9.6 +/− 0.9 Hz, right hemisphere: 9.7 +/− 0.9). The correlational and instantaneous phase difference of the envelopes were also comparable to the sensor data (r_Env,Source_ = 0.37 ± 0.19, Φ_Env,Source_ = 3.2° ± 23.9°). The phase offset of the IAF signal in source space trended towards anti-phase, however, inter-subject variability was much larger than in sensor space (Φ_IAF_ = −138.6° ± 76.6°) such that the signals were uncorrelated when averaged across subjects (r_IAF_,_Source_ = 0.00 ± 0.48). This further emphasizes that the low frequency envelope rather than the IAF signals exhibit synchronous oscillations between hemispheres.

We then designed the tACS waveforms with the IAF of each subject as the main carrier frequency, while the amplitude was modulated with a fluctuating 1 Hz envelope (maximal Intensity = 1.5mA peak-to-peak, Amplitude modulation = 0.5 of maximal intensity). We employed two stimulation regimes: In the *phase-synchronized* condition, the relative phase of IAF was 0° (Φ_IAF_ = 0°) and the relative phase of the 1 Hz envelope was 180° (Φ_Env_ = 180°), whereas this was reversed in the *power-synchronized* condition Φ_Env_ = 0° and Φ_IAF_ = 180°; Figure 1D). Note that power-synchronized tACS approximates the interhemispheric coupling measured with EEG at rest (correlated envelopes, anti-/uncorrelated IAF signals). Next we modeled the electric field to assess whether our two stimulation paradigms caused comparable effects in sensorimotor cortex. We found that both the amplitude of the alpha oscillations and the amplitude of the power-envelope were similar between the two stimulation paradigms over sensorimotor cortices (Figure 2), such that the relative phase relationship of the IAF/power-envelope between the left and right hemisphere represent the major difference between the two stimulation protocols when analyzed within SM1. Note, however, that the three-electrode setup did not allow a perfect matching of the electric field distribution across the whole brain. In particular the phase-synchronized stimulation also produces a strong electric field over occipital areas, which follows the carrier frequency but is no longer modulated by the envelope. Most importantly the power-synchronized stimulation paradigm produces an in-phase fluctuation of the IAF envelope in the sensorimotor network, whereas phase-synchronized stimulation produces an anti-phase fluctuation of the power-envelope in the sensorimotor system.

### tACS effects on the sensorimotor RSN

The same 20 subjects participated in a second session where we applied tACS using a three-electrode setup (Figure 1D, similar to (Polanía et al., 2012)) to the left and right SM1 inside an MR scanner to modulate rs-fMRI connectivity. In total, we acquired 5 resting-state scans for each subject (Figure 1D; 7min, eyes-open, 3 minutes break between blocks). After measuring baseline connectivity (1^st^ run), either phase-synchronized or power-synchronized tACS was applied in the 2^nd^ and 4^th^ run respectively (order counterbalanced across subjects), and stimulation aftereffects were measured in runs 3 and 5 (POST-phase-synchronized and POSTpower-synchronized). Subjects were aware that they would be stimulated and 16 subjects reported feeling a slight tingling in the scanner, but they were not able to distinguish between the two stimulation regimes as established prior to the rs-fMRI scans via verbal report.

Resting state networks (RSNs) were spatially identified by group independent component analysis (ICA) based on the whole dataset and we further analyzed eight pre-selected RSNs of interest for further analysis (Figure 3): the default-mode network (frontal and dorsal part, Figure 3 red), a sensorimotor network (Figure 3 dark blue), a premotor-network, the striatum (Figure 3, blue-green), a lateral motor network (Figure 3, green), a superior parietal network (Figure 3, light blue) and a visual network (Figure 3, yellow). We then compared the influence of the two tACS regimes on the different RSNs using a dual regression approach (Stagg et al., 2014).

**Figure 3.**
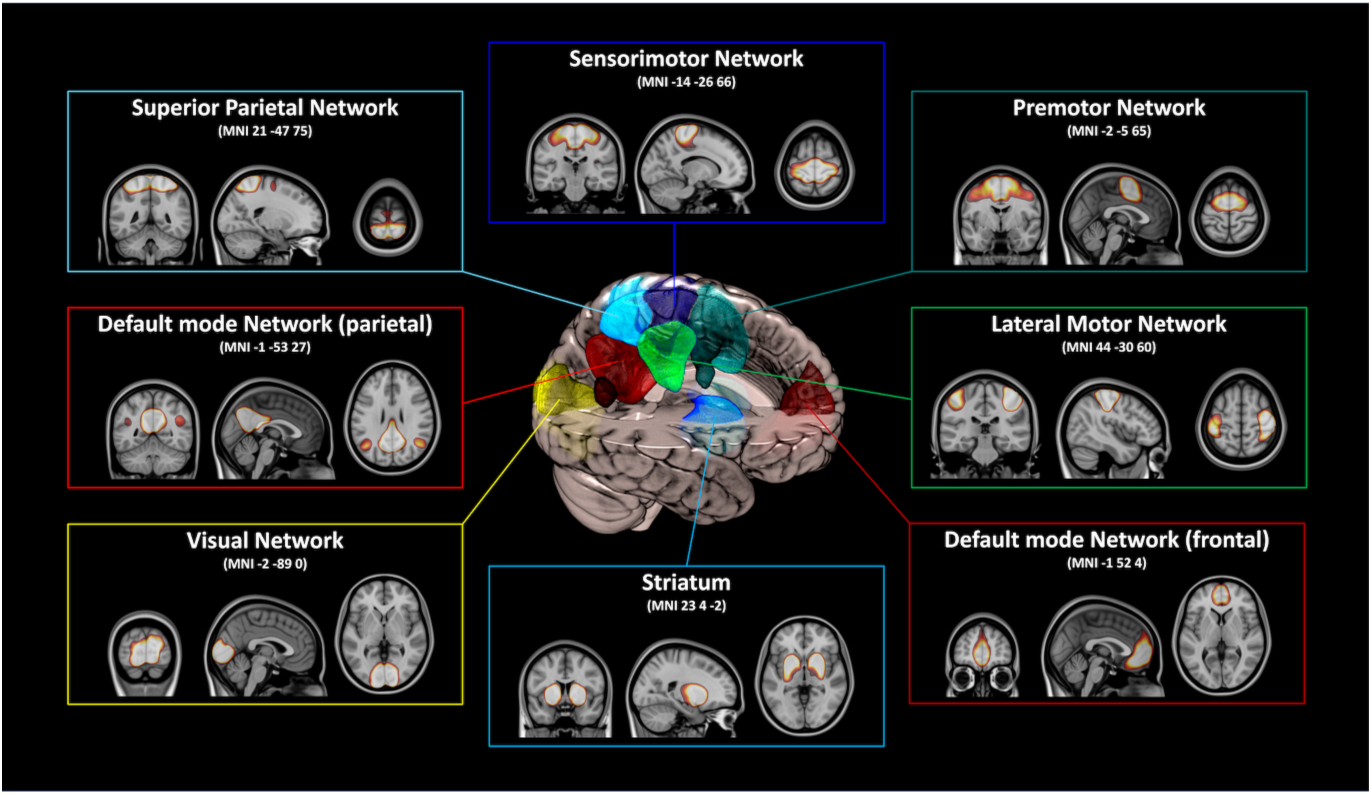
Resting state networks of interest. Based on the MELODIC group-ICA, eight resting-state networks were selected for further analyses: A sensorimotor network (blue), a premotor network (blue-green), a more lateral motor network (light green), the default-mode network split into a frontal (dark red) and parietal part (light red), the striatum (light blue, bottom), a visual network (yellow) and a superior parietal network (light blue, top).

This approach yields a surrogate measure of synchronicity or coupling strength within the RSN under investigation.

Across all RSNs of interest, only the sensorimotor network changed its coupling strength significantly when tACS was applied (main effect of stimulation type F(4,72) = 4.462, p = 0.003 in a linear mixed effects model including all five rs-fMRI runs). No other RSNs exhibited significant modulation effects (although the pre-motor network and the striatum approached significance, which is not surprising since these RSNs are strongly interconnected with motor areas; premotor network: F(4,72) = 2.484, p = 0.051; striatum: F(4,72) = 2.102, p = 0.089).

Next we calculated a normalized index of network strength within the sensorimotor RSN to determine changes relative to the first baseline rs-fMRI scan (linear mixed effects model, 4 rs-fMRI runs normalized to baseline; Figure 4). Again, we found a main effect of stimulation type only for the sensorimotor system (F(3,57)=4.402, p=0.007). Power-synchronized tACS increased connectivity strength by 25% ± 9% (mean ± s.e.m.), which was significantly stronger than the effect of Phase-synchronized tACS (8% ± 7% increase; p = 0.037, Cohen’s d = 0.49). Moreover, we found *after-effects* of the power-synchronized tACS protocol on RSN connectivity (34% ± 8%; compared to baseline) that were significantly larger than after-effects (12 ± 6%, p=0.01, Cohen’s d = 0.4) or acute effects (8% ± 7%) caused by phase-synchronized tACS (p = 0.002, Cohens d=0.62). Note that analogous results were revealed in a control analysis where the spatial topography of the RSNs was determined from an independent data set, showing that the effect was robust to variations in RSN anatomy.

**Figure 4.**
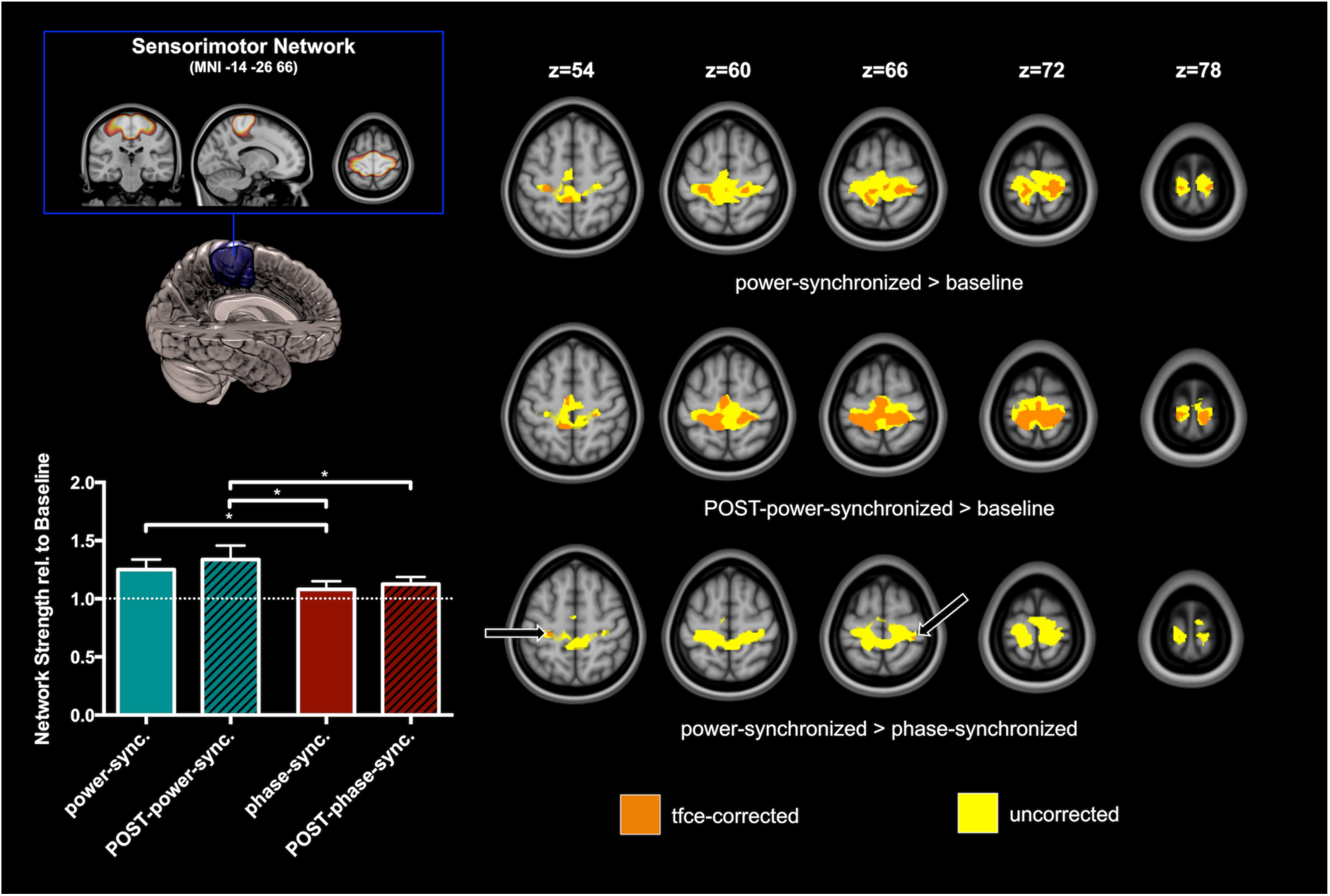
Increase of network strength of the sensorimotor network during power-synchronized tACS compared to baseline and phase-synchronized tACS (control). (Left) Averaged normalized scores of network strength identified by dual regression during and after power-synchronized (blue-green) and phase-synchronized tACS (dark red) revealing an increase of 25% in network strength during power-synchronized tACS compared to baseline and a significant increase (20%; p = 0.037) when compared to phase-synchronized tACS (control). The increase was still significant when comparing the two after-effects (POST-power-synchronized and POST-phase-synchronized, p = 0.010; *p < 0.05). (Right) Corresponding voxelwise contrasts revealing an increase of network strength around the central sulcus in between the active electrodes during and after power-synchronized tACS compared to baseline and phase-synchronized tACS, and a general increase during and after power-synchronized tACS when compared to baseline (all images: p_tfce-corr._ < 0.05).

Importantly, after normalization to the baseline scan the sensorimotor network was again the only network across all RSNs of interest that showed a significant main effect. Only the pre-motor network, but no longer the striatum, showed a trend towards significance (Figure 5).

**Figure 5.**
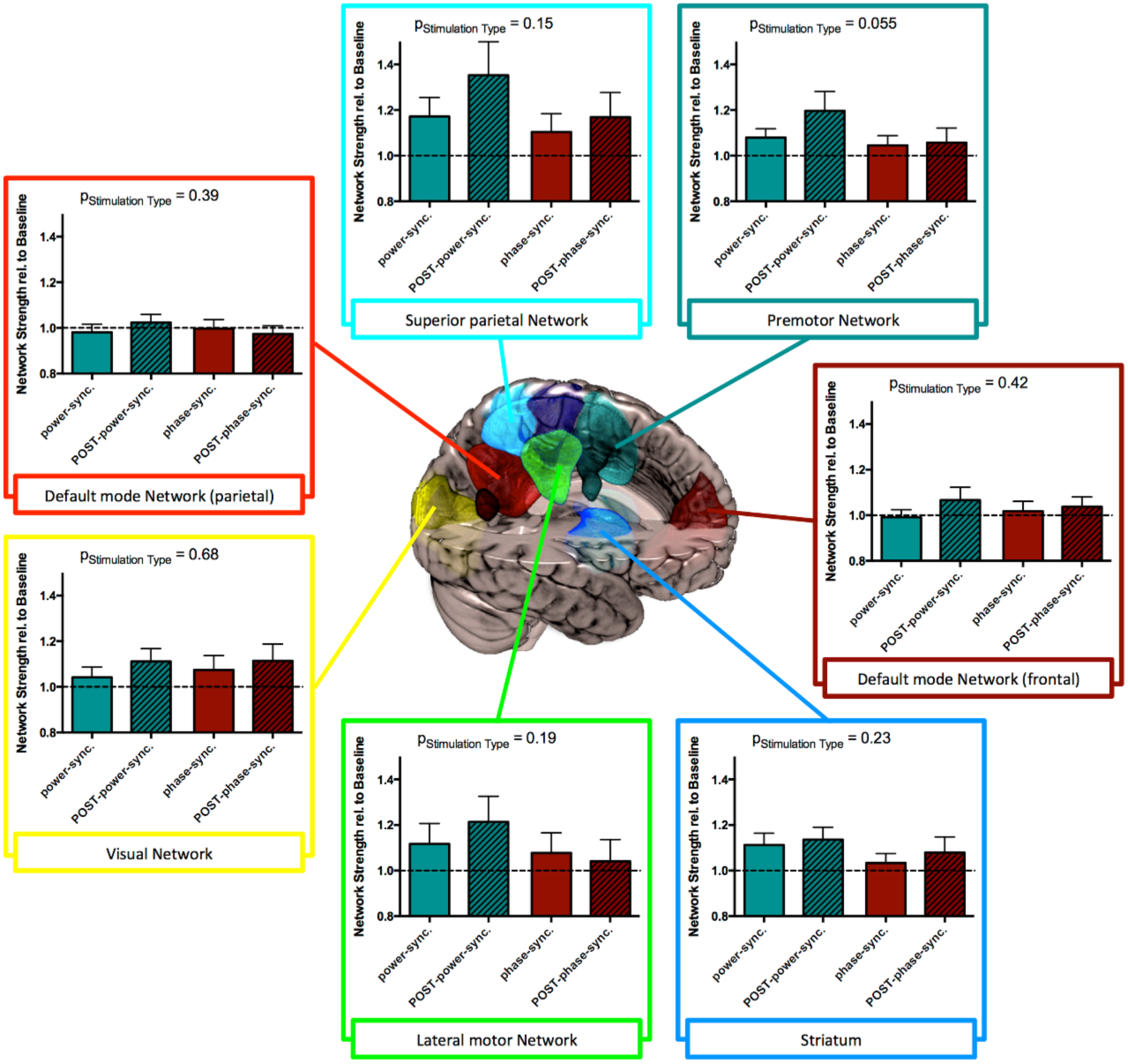
Results of other resting state networks of interest. We did not find a main effect of stimulation type for any RSN other than the sensorimotor network. However, there was a similar trend (i.e. power-synchronized tACS increasing network connectivity vs. phase-synchronized tACS not increasing network connectivity) observable in the striatum, the premotor network, the lateral motor network and the superior parietal network. Although the effect did not reach statistical significance, the general trend fits the observed changes in the sensorimotor network since all of these networks are either associated with motor functions, and are therefore connected to the sensorimotor network (premotor network, striatum), or in between/under the stimulation electrodes (superior parietal network, lateral motor network). By contrast, we observed no difference between stimulation paradigms for the default-mode network (parietal & frontal) and the visual network which rules out that the above described effects are purely driven by current injection itself or current related artifacts. p_Stimulation Type_ given for normalized data.

Since we observed after-effects following both types of stimulation we also calculated how tACS changed the network strength relative to the resting-state run immediately before stimulation (Figure 6). For power-synchronized stimulation we found similar increases during (14% +/− 5% mean +/− s.e.m.) and after stimulation (22% +/− 10%). For phase-synchronized tACS we no longer observed an increase in connectivity during (−4% +/− 6%) or after stimulation (2% +/− 6%). Statistics still revealed a main effect of stimulation (LMEM, F(3,57) = 3.128, p=0.033) with significant post-hoc differences between the after-effects of the two stimulation types (p=0.032) and a strong trend for the acute stimulation effects (p=0.067).

**Figure 6.**
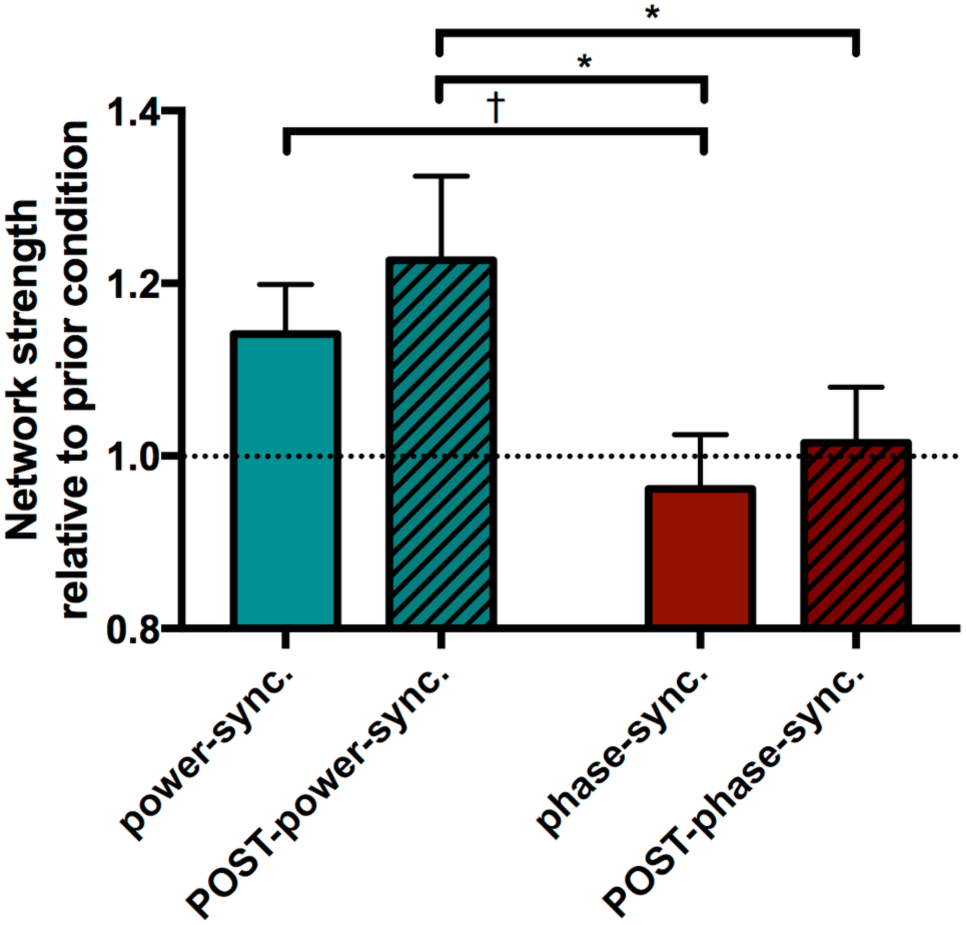
tACS induced changes in network strength relative to the previous condition. Note that only power-synchronized stimulation signals cause a relative increase of connectivity strength while phase-synchronized stimulation (i.e. power envelopes are in anti-phase) has only a minor effect on connectivity. The bars show the mean and standard error (^*^p<0.05, ^†^p<0.1).

We then analyzed order effects of the stimulation conditions, i.e. we split the group of participants into those where the power-synchronized condition was applied first (i.e. following the baseline measurements) and those where the power-synchronized stimulation was applied second (i.e. on top of the after-effects induced by the phase-synchronized stimulation (Figure 7). These results suggest that power-synchronized stimulation had a synchronizing effect irrespective of order (16% +/− 5% when applied as the first stimulation, and 12% +/− 6% when applied as the second stimulation). The phase-synchronized condition, however, only had a slight synchronizing effect when applied first (5% +/− 7%), but a desynchronizing effect when applied second (−13% +/− 7%).

**Figure 7.**
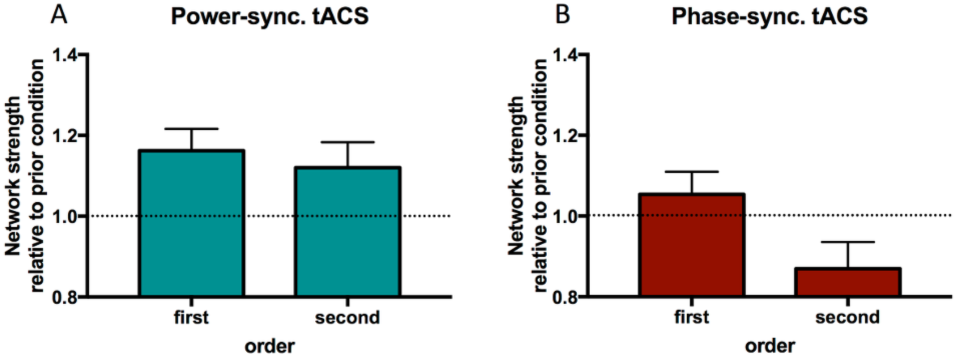
Effects of power-synchronized (left) and phase-synchronized stimulation (right) on network connectivity strength when participants were split depending on stimulation order. The data are normalized to the rest period immediately before stimulation. Note that power-synchronized stimulation has a synchronizing effect on the sensorimotor network irrespective of whether it followed the baseline condition or was applied on top of the after-effects caused by the phase-synchronized stimulation. Phase-synchronized stimulation has a slight synchronizing effect when applied immediately after baseline, but a de-synchronizing effect when the network state had already been modulated by power-synchronized tACS.

Next, we performed a voxelwise analysis within the sensorimotor network to localize the effect previously observed at the level of the averaged network strength.

Contrasting voxelwise RSN strength during power-synchronized tACS versus the baseline scan revealed a large cluster of voxels around the central sulcus, extending into the left middle cingulate cortex, left post central gyrus and the right pre- and postcentral gyrus. (Figure 4, power-synchronized > baseline, p_tfce-corr._ < 0.05, threshold-free cluster enhancement corrected). The size of this cluster increased even more when looking at after-effects by comparing POST-power-synchronized to baseline (paired t-test POST-power-synchronized > baseline, p_tfce-corr._ < 0.05). By contrast, no voxel survived statistical thresholding when comparing phase-synchronized tACS or its after-effects to baseline (p_tfce-corr._ > 0.05 for all voxels). The direct comparison of connectivity measured during power-synchronized versus phase-synchronized tACS revealed only two very small clusters close to the central sulcus in the left pre- and postcentral gyrus and the right postcentral gyrus (power-synchronized > phase-synchronized, p_tfce-corr._ < 0.05). Note that according to our simulation, power-synchronized stimulation produces an electric field that peaks over the post-central gyrus and stretches all along the central fissure from parietal to premotor regions (Figure 2B). We would therefore expect the strongest effect for power-synchronized stimulation over post-central regions, which is in line with the voxel-vise analysis.

Our setup used a single return electrode over visual cortex that was the same size as the two stimulation electrodes. It is possible that this electrode placement might have induced subliminal phosphenes, especially during phase-synchronized stimulation, where the electric field is estimated to be almost twice as strong over occipital areas. Even though none of our subjects reported awareness of visual stimuli in the scanner when de-briefed, possibly due to a relatively weak stimulation intensity (max. amplitude 750 μA) in combination with the dimly lit scanner room, we performed a control analysis focused on the visual RSN. However, we found no statistical evidence that the visual RSN may have been modulated by either power-synchronized or phase-synchronized tACS stimulation (Figure 5). This makes it very unlikely that the effects reported above were artifacts caused by visual flicker, a common side effect of tACS. This argument is further supported by the fact that the stronger rs-fMRI coupling within the sensorimotor RSN also persisted during the after-effect runs (POST runs, without tACS stimulation).

Even though the electrical field induced by phase-synchronized and power-synchronized stimulation was similar within the sensorimotor RSN (Figure 2), it is not possible to match the induced electrical fields across the whole brain with a three-electrode setup. In particular, the simulation revealed that phase-synchronized tACS affected parietal and occipital regions by entraining synchronous activity across hemispheres at the IAF. However, this did not result in higher RSN connectivity suggesting that phase-synchronized stimulation was less effective in influencing rs-fMRI connectivity than power-synchronized stimulation. This was further confirmed by a control analysis of a parietal network (extending bilaterally from superior parietal cortex and middle occipital gyrus, and including a small cluster in right inferior frontal gyrus) which revealed no effect of stimulation type (F(3,57)=0.345, p=0.793).

Finally, we found no stimulation specific effect for the default-mode network (parietal& frontal), the superior parietal network or the lateral motor network, which rules out that the reported effects are purely driven by current injection itself or by artifacts possibly related to the current (Figure 5). However, the superior parietal, premotor and lateral motor network showed a similar response pattern as the sensorimotor network. We therefore ran an analysis on a full model (LMEM) incorporating all networks of interest and stimulation types. We found a significant main effect of stimulation type (F(3,513) = 4.979, p = 0.002) supporting our main claim that power-synchronized tACS is more effective for increasing rs-fMRI connectivity than phase-synchronized tACS. We also found a significant main effect of network (F(6,513) = 5.176, p<0.001) but the stimulation type*network interaction was not significant (F(18,513) = 0.646, p=0.863), most likely because the electrical field spread widely (see Figure 2B) and because intersubject variability was large. We also calculated Cohen’s d effect sizes between the power-synchronized versus phase-synchronized conditions for each of the networks and found a medium effect size only for the sensorimotor network (Cohen’s d=0.464). The next largest effect was in striatum (Cohen’s d=0.36), while the effect sizes for all other networks were small (Cohen’s d= −0.152 to 0.17). Thus, power-synchronized tACS had the strongest effect on the sensorimotor network even though other networks were also mildly affected.

### Relationship between EEG connectivity and rs-fMRI connectivity within the sensorimotor RSN

Next, we analyzed the effect of power-synchronized versus phase-synchronized tACS for each participant and found high inter-individual variability (Figure 8). To understand the possible origins of this variability we investigated whether there is a relationship between interhemispheric connectivity measurements derived from the EEG data and the modulatory effect of the power-synchronized or phase-synchronized tACS on rs-fMRI connectivity in the sensorimotor RSN. We found a negative correlation between the r_Env_ (Figure 1B, blue) and the percentage increase in rs-fMRI connectivity strength within the sensorimotor network caused by power-synchronized tACS (Figure 9A; spearmans rho = −0.481, p = 0.032). Note, however that this correlation does not survive Bonferroni correction. Nonetheless this indicates that subjects for whom the initial coupling of the IAF’s power envelope was low tended to show a greater synchronization effect in response to power-synchronized stimulation. Moreover, we observed that those subjects who exhibited a strong natural interhemispheric coupling of IAF (i.e high r_IAF_) tended to respond stronger to power-synchronized tACS than those with a weak IAF coupling (Figure 1B, red). Even though this association just failed to reach statistical significance (Figure 9B; Spearman’s rho = 0.432, p = 0.057), it further suggests that rs-fMRI connectivity mostly reflects synchronous activity of the power-envelopes.

**Figure 8.**
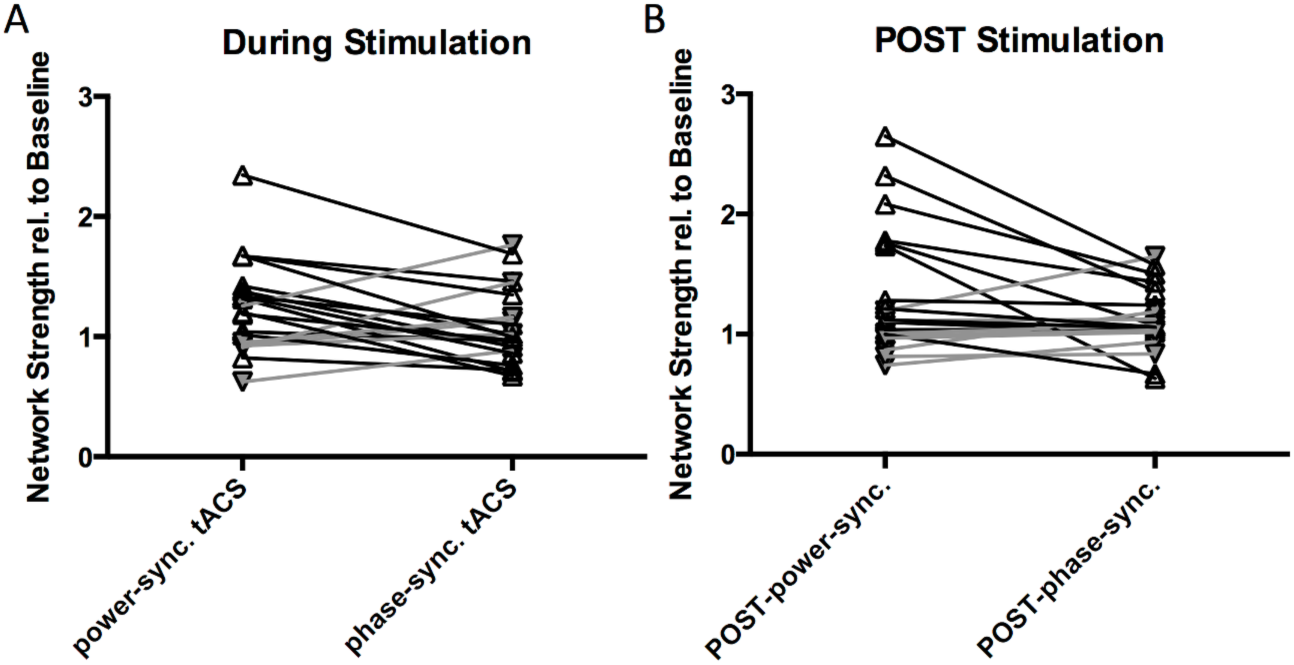
Individual subject data. (Left) Normalized network strength of the sensorimotor network during power-synchronized compared to phase-synchronized tACS and (Right) during POST-power-synchronized and POST-phase-synchronized for each subject revealing large intersubject variability. Subjects who do not follow the general trend (power-synchronized tACS > phase-synchronized tACS) are depicted in gray.

**Figure 9.**
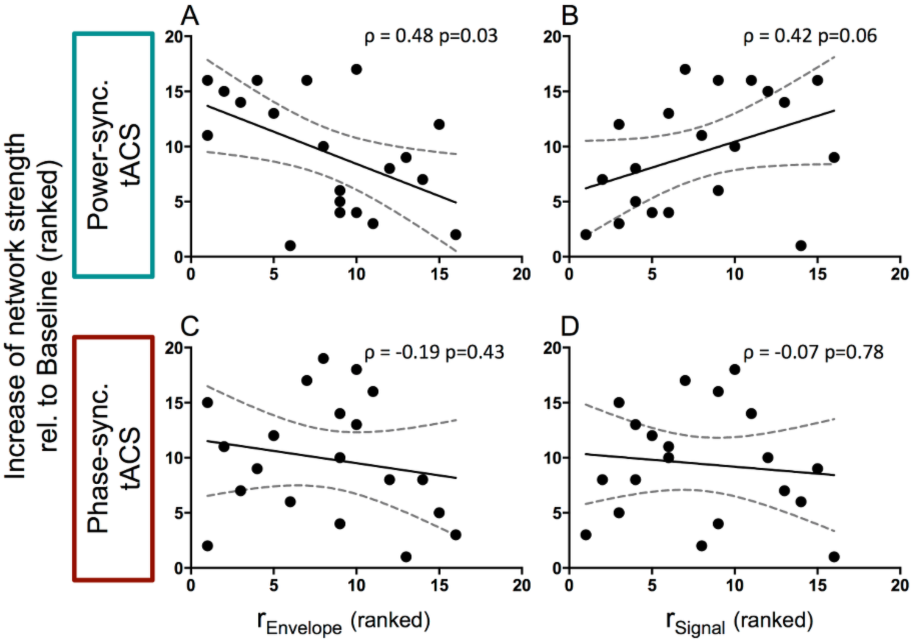
Correlations between EEG signal properties and percentage increase of network strength during power-synchronized and phase-synchronized tACS. (A) Correlation between increase of network strength during power-synchronized tACS and IAF envelope correlation between electrode C3 and C4 (r_Envelope_) revealing a negative association between the IAF envelope over sensorimotor cortex in the sensor space and efficacy of tACS stimulation (rho = −0.48 p = 0.03). (B) Positive correlation of increase of network strength during power-synchronized tACS and IAF signal correlation between electrode C3 and C4 (r_Signal_) (rho = 0.42 p = 0.06). (C,D) No such relationship was visible between the change in network strength during phase-synchronized tACS and IAF envelope or signal between C3 and C4 (IAF envelope: rho = −0.19, p = 0.43 IAF signal: rho = −0.07, p = 0.78). Note that EEG and fMRI measurements were performed in two separate sessions.

No such correlations were found when r_IAF_ and r_Env_ were related to the percentage increase in rs-fMRI connectivity strength caused by phase-synchronized tACS (Figure 9C,D).

Since phase-synchronized tACS could potentially influence alpha activity in occipital cortex, we ran control analyses to test whether there was a correlation between increase in network strength and occipital alpha power or between increase in network strength and the difference between sensorimotor and occipital alpha frequency. Neither of these correlations were significant (p>= 0.12).

## Discussion

The present results show that entraining the alpha-power envelopes of remote yet connected areas in a synchronized fashion strengthens rs-fMRI connectivity within the targeted RSN. Increased connectivity strength was consistently observed irrespective of whether the effect of power-synchronized tACS was compared to baseline or connectivity strength immediately prior to stimulation. This increase in connectivity outlasts the actual tACS stimulation period, suggesting some form of adaptation within the targeted neural circuits (Neuling et al., 2013; Vossen et al., 2014). Additionally, we demonstrated that the power-synchronized tACS regime was more effective in modulating rs-fMRI connectivity when applied in individuals with weak electrophysiological interhemispheric coupling (i.e., less synchronized IAF power-envelopes). Importantly, only small, non-significant effects were observed when the tACS signals phase-synchronized the IAF rhythm while the power-envelopes oscillated in anti-phase. Together, these findings provide causal evidence that power-synchronization is an effective mechanism for linking oscillatory neural activity across different temporal scales. In particular, they demonstrate how the mechanism of cross-frequency coupling enables fast oscillatory rhythms to influence ultra-slow oscillations measured by inter-areal BOLD connectivity, an idea that was until now only supported by correlative evidence.

Finally, our results indicate that long-range connectivity can be modulated with tACS if the stimulation signals are appropriately designed. This might not only open new possibilities for experimental research, but also lead to new therapeutic applications in diseases where resting-state connectivity is diminished, as for example in Autism or Alzheimer’s disease (Greicius, 2008; Alaerts et al., 2014).

Here we used the alpha rhythm as a carrier frequency because it can be entrained by tACS (Zaehle et al., 2010; Neuling et al., 2013) and previous research revealed a systematic link between alpha power and the BOLD signal (Mantini et al., 2007; Ritter et al., 2009; Brookes et al., 2011; Hipp et al., 2012; Wang et al., 2012). However, power-synchronization most likely represents a general mechanism which might be independent of the carrier frequency. Moreover, it is likely that rs-fMRI connectivity can also be modulated by directly synchronizing oscillations at 1 Hz or below (e.g. via optogenetic manipulations).

One might be surprised that applying phase-synchronized tACS, during which the power-envelopes oscillated in anti-phase, did not *de*synchronize the BOLD signal. However, synchronized IAF signals themselves might have a stabilizing effect if BOLD connectivity is relatively weak (Wang et al., 2012) as suggested by Figure 6, which shows that phase-synchronized tACS enhanced connectivity when the brain was in its typical connectivity state, as measured during the baseline scan (Figure 7. right, condition first). However, if connectivity was enhanced, phase-synchronized tACS tended to reduce rs-fMRI connectivity (Figure 7, right, condition second). By contrast, power-synchronized tACS consistently increased rs-fMRI connectivity measures irrespective of the prior state (Figure 7, left).

In addition to the acute effects of our stimulation paradigm, we also found significant increases in the resting-state run immediately after power-synchronized tACS.

A recent study reported changes in alpha-power for up to 40 minutes after 10 minutes of tACS stimulation (Neuling et al., 2013). Currently, it is hypothesized that these prolonged tACS stimulation effects result from spike-timing dependent plasticity (STDP), or from long term potentiation/depression (LTP/LTD (Vossen et al., 2014)) causing an extended period of increased long-range connectivity as observed during the POST-power-synchronized runs. Even though our study cannot speak to the underlying mechanism it is promising from a therapeutic perspective that a relatively short stimulation period of 7 minutes causes longer lasting after-effects in modulating interhemispheric coupling strength. It would be interesting to test whether this after-effect, which we measured at rest, would also be beneficial in the context of performing a task.

Our results suggest that there is an inverted relationship between the correlation of the EEG signal and the effect seen during power-synchronized tACS (i.e., low EEG correlation leads to a strong increase in rs-fMRI connectivity during power-synchronized tACS). No such relationship was observed with the phase-synchronized tACS condition. Two potential explanations for this observation are that entrainment is more efficient if the intrinsic interhemispheric connectivity of the IAF power envelopes is naturally weak, or that tACS entrainment does not yield additional benefits if the network is already highly synchronized.

### Distribution of the electric field

One problem concerning non-invasive brain stimulation studies is the design of an appropriate control condition. Here we asked whether synchronizing 1 Hz oscillations across the left and right sensorimotor cortices via power-synchronized tACS increased connectivity within the sensorimotor network. We compared this to a control condition which used the same principle tACS signals but the relative phase was shifted such that the IAF rhythms were synchronized while the power envelopes were in anti-phase. Although the electric field strength is comparable between both tACS conditions for the sensorimotor network, the overall distribution of the electric field across the rest of the brain and the direction of the induced current differed. In particular, phase-synchronized stimulation produces an electric field shifted more towards occipital areas resulting in IAF stimulation that also had a strong effect on the primary visual cortex. Nevertheless, phase-synchronized tACS was rather ineffective for increasing connectivity within any of these RSNs even though the modulation of IAF amplitude was relatively strong (up to 0.4 V/m peak-to-peak). This suggests that entrainment of rs-fMRI connectivity seems not to depend on the induced field strength alone, but also on the entrained frequency bands and their phase relationships. Our study suggests that the power-synchronized stimulation regime is suitable for increasing rs-fMRI connectivity within RSNs and it is possible that both the 0° phase-shift of the power-envelope and the 180° phase-shift of the IAF are necessary for inducing this effect. In particular, it is likely that the carrier frequency (i.e. IAF) is necessary to entrain neuronal populations of cortex, but that the slow fluctuating envelope is necessary to synchronize fluctuations in the BOLD signal, which is measured by rs-fMRI connectivity metrics. This interpretation would explain why neither 1 Hz nor IAF stimulation alone were suitable for enhancing rs-fMRI (Vosskuhl et al., 2015). Moreover, it is in line with previous research suggesting that low frequency components of the LFP (Lu et al., 2014) as well as low-frequency power-envelopes of EEG/MEG oscillations are particularly strongly associated with BOLD fluctuations (Mantini et al., 2007; Brookes et al., 2011; Hipp et al., 2012).

## Conclusion

Most studies in humans relied on correlational approaches to investigate the relationship between fMRI measurements and electrophysiology since invasive electrical stimulation of the brain (i.e., direct electrical stimulation of cortex or deep brain stimulation) is usually restricted to clinical settings (Mandonnet et al., 2010; Bronstein et al., 2011). Compared to these methods tACS has the advantage of being non-invasive and easy to apply. Here we showed that if the waveforms are well designed, tACS is an appropriate tool for modulating the strength of resting-state networks. In particular, we provided causal evidence that power-synchronization is effective for linking oscillatory neural activity across different temporal scales. This could open new opportunities for human fMRI research and new interventional approaches for modulating long-range connectivity of the human brain at rest. Potential therapeutic applications could be, for example, tACS-based modulation in patients with diminished resting-state connectivity within specific circuits, as for example in Autism (Alaerts et al., 2014) and Alzheimer’s disease (Greicius, 2008).

## Acknowledgements

The authors would like to thank Karl Treiber for scanning assistance and Daniel Woolley for feedback on the manuscript.

The research was supported by the Swiss National Science Foundation (320030_149561 & 320030_146531); the NCCR Affective Sciences; FWO Flanders (G.0401.12) and the Seventh Framework Programme European Commission (PCIG12-2012-334039).

